# Novel methodologies for the treatment of type 1 diabetes mellitus using *in silico* tools

**DOI:** 10.1101/2025.05.30.657014

**Authors:** Filipe Ricardo Carvalho, Paulo Jorge Gavaia

**Affiliations:** Faculty of Medicine and Biomedical Sciences, University of Algarve, Faro, Portugal; Centre of Marine Sciences (CCMAR/CIMAR LA), University of Algarve, Faro, Portugal Universidade do Algarve - Campus de Gambelas, 8005-139 Faro, Portugal

## Abstract

Epidemiologic studies reveal that the fight against diabetes is being lost, so to overcome this serious health problem every strategy should be taken seriously. Using *in silico* data we discuss the therapeutic potential for Diabetes Mellitus type 1 of two possible unstudied strategies; (I) if *INSULIN* paralog, *INSULIN-INSULIN GROWTH FACTOR 2* can be a regulator of glucose in humans, and (II) if extrapancreatic expression of *INSULIN* and *INSULIN- INSULIN GROWTH FACTOR 2* has any relevance in glucose regulation. To address this hypothesis, we access to the collection of protein sequences and gene expression from INSULIN and paralogues in EGB, NCBI and EBI. Phylogenetic and molecular evolutionary analyses were conducted using MEGA software, conserved protein motifs were compared using algorithm MUSCLE and presenting it graphically using CLC Genomics Workbench, transcripts and gene expression from transcriptome and microarray data from different experiments of insulin and paralogues were extracted and graphicaly presented using CIRCOS. Phylogenetic analysis revealed that humans, like most of all mammals, have *INSULIN* and *INSULIN-INSULIN GROWTH FACTOR 2* with the exception of murinae among mammals that share *Insulin 1* and *Insulin 2*. Both *Insulin 1 and 2* can regulate glucose since both proteins are highly similar and with a redundant function, as confirmed by *in vivo* experiments showing that both knockout *Insulin 1* or *2* are normoglycemic. Human INSULIN- INSULIN GROWTH FACTOR 2 lacks motif 3 of the INSULIN signature, and part of the predicted protein structure does not present any known domain, making the prediction of a possible function for this gene inconclusive. Extrapancreatic expression of insulin has been observed in diabetic mice with contradictory effects, but available gene expression data in repository data banks also suggests that extrapancreatic expression of insulin occurs in human, mouse and zebrafish under basal condition. Gene expression DataSets demonstrate that extrapancreatic expression of INSULIN occurs under basal condition. Further investigation should be made, to understand if extrapancreatic expression of insulin can have a role in glucose homeostasis, as this could represent a complementary strategy in diabetes treatment.

## INTRODUCTION

Since the discovery of insulin in 1921 ^[1]^, patients with type 1 Diabetes Mellitus (T1DM) passed from a fatal prognostic to chronic patients, thanks to insulin replacement therapy. Although this treatment is the most widely peptide or protein based medicine used in the world ^[2]^, it was shown not to be totally efficient in treating all insulin-dependent diabetics, leading to morbidity later in life. Mouse has been the most widely animal model used for the study of diabetes ^[3]^, but recently, protocols for the induction of diabetes and transgenic lines in zebrafish have been developed ^[4,5]^. The canonical gene of insulin in mouse is *Ins2* and in zebrafish is *insa* while in human it is *INS*

(Human insulin gene) and all sharing a high percentage of identity. In these three species we can find a paralog for insulin gene, being in mouse *Ins1*, in zebrafish insb and in human *INS-IGF2*. In human it has been identified a paralog of the insulin gene, *INS-IGF2* ^[6]^, which is a conjoined gene sharing *INS* gene in the 5’ region and the *IGF2* gene in the 3’ region. *INS-IGF2* is a parental imprinting gene that was first identified as being expressed both in the eye and pancreas ^[6]^ and a possible autoantigen like *INS* in T1DM ^[7]^. In mouse, the two genes encoding for insulin, *Ins1* and *Ins2*, are thought to have duplicated because of an infection in the genetic material by a RNA virus originating *Ins1* (retrogene). Investigations done with mouse *Ins1* or *Ins2* knockouts (KO) revealed that *Ins1* alone can regulate glucose homeostasis, although in some cases it cannot prevent type 1 diabetes mellitus (T1DM). Duvillie et al. ^[8]^ showed that double *Ins1* and *Ins2* KO mice died prematurely, but mice lacking just one insulin gene were viable and fertile ^[8]^. Leroux et al. ^[9]^, working with the same mutant lines observed, not only, that both *Ins1* and *Ins2* KO mice were normoglycemic, but that Ins2 KO mice had a dramatic increase in the expression of *ins1* accompanied with an increase in β-cell mass ^[9]^. Babaya et al. ^[10]^, showed that *Ins2* KO nonobese diabetic (NOD) male mice, with just one *ins1* allele, NOD^ins1+/-,ins2-/-^, developed diabetes at around 10 weeks of age, but males with both ins1 alleles NOD^ins1+/+,ins2-/-^ and females from the two mutant lines only presented signs of diabetes later in life ^[10]^.

In zebrafish the existence of two insulin genes (*insa* and *insb*) is likely due to the genome duplication event that occurred in the stem lineage of teleost fishes ^[11]^. Papasani et al. ^[12]^, identified the second insulin gene in zebrafish and observed expression of *insb* both in pancreas and brain and suggested that it may regulate both autocrine/paracrine and endocrine systems via both *insulin receptor a (insra) and insulin receptor b insrb* (14).

In human extrapancreatic expression of insulin was first identified in brain ^[13]^ and then in thymus ^[14]^, has a way to the immune system recognize insulin, avoiding autoimmunity and β-cell destruction. Later Kojima et al showed the presence of cells that stained positive for insulin RNA in the liver, adipose tissue and bone marrow in several diabetic mice models but not in nondiabetic mice ^[15]^. However, this insulin expressing cells had no impact on hyperglycemia in diabetic mice. Later, Kojima et al. ^[16]^ demonstrated, that beside not having any impact in regulating glucose, this Proins/TNF-α- expressing cells had their origin in bone marrow and then migrated to several parts of the body, initiating diabetic neuropathy ^[17]^. Cunha et al ^[18]^, in their experiments with diabetic mice treated with streptozotocin, observed beneficial effects in the secretion of insulin by the tear film of the eye and locally synthesized in the lachrymal gland ^[18]^. So at the present moment Human and animal studies, points in the direction that extrapancreatic occurs in the different species, but it is not clear what are the tissues that express INS, in what situations and most importantly can it be in countervailable quantities to replace pancreatic INS.

### Objectives

Filling the gap in knowledge, of how this phenomenon occurs and if it can represent a novel therapy for DM seems to be a issue that needs to be far-fetched and to the best of our knowledge few^[19]^, in recent years, have been focused in this subject. With the objective to understand if extrapancreatic expression occurs, using *in silico* analyses, we proposed to understand; 1) if INS-IGF2 can be a regulator of glucose homeostasis and 2) if extrapancreatic expression of both INS and INS-IGF2 in Humans can occur mimicking pancreatic INS action.

## METHODS

### Study design

#### Phylogenetic tree construction

To better understand the evolution of insulin genes we conducted a phylogenetic analysis of *INS, INS-IGF2, Ins1, Ins2, insa* and *insb* together from available sequences human, chimpanzee, orangutan, gibbon, marmoset, otolemur, tree shrew, mouse lemur, squirrel, lesser jerboa, chinese hamster, rat, mouse, ferret, cat, panda, dolphin, cow, elephant, manatee, walrus, microbat, guine pig, hyrax, marmoset, coelacanth, frog, anole lizard, green sea turtle, duck, chicken, turkey, rock dove, collared flycatcher, zebra finch, ground tit, medium ground finch, white throated sparrow, tilapia, japonese pufferfish, green spotted puffer, zebrafish, medaka, platyfish, greater amberjack, stickleback and ciona intestinalis. Collection of protein sequences from insulin and paralogues was obtained in Ensembl Genome Browser (EGB) ^[20]^ and protein database of National Center of Biotechnology Information (NCBI) ^[21]^.

We collected 32 INS, 11 INS-IGF2, 2 ins1, 2 ins2, 7 insa and 5 insb protein sequences from the different species analyzed (Figure 2). For multiple sequence alignment we used the algorithm Multiple Sequence Comparison by Log- Expectation (MUSCLE) ^[22]^. Based on the sequence alignment results, the phylogenetic tree reconstruction was made using the Maximum-likelihood statistical method. Test of phylogeny was done using the bootstrap method and the number of replications was 1000. Phylogenetic and molecular evolutionary analyses were conducted using MEGA version 6 ^[23]^. The insulin like 1 precursor of 275 amino acids sequence of ciona intestinalis was used as an outgroup in the phylogenetic tree.

#### Graphical representation of conserved motifs

To determine percentage of identity between paralogs we aligned both proteins sequences for human, mouse and zebrafish species using algorithm MUSCLE and presenting it graphically using CLC Genomics Workbench v6.5 (CLC bio, Aaarhus, Denmark). Also, we applied *INS, INS-IGF2, Ins1, Ins2 insa* and *insb* to InterProScan database ^[24]^ to identify insulin signatures and motifs present in protein sequences.

#### Transcripts and gene expression

To understand the pattern of expression of insulin genes in different tissues we gathered information available in data bases that provide transcriptome and microarray results from different experiments. Tissue gene expression pattern from *insa* (Dr.75811), *insb* (Dr.87912) and *Ins2* (Mm.4946) were obtained from UniGene data base ^[25]^ and *INS* ^[26]^, *INS-IGF2* ^[26]^ and *Ins1* ^[27]^ from Expression Atlas ^[28]^. No statistical analyses was performed, as all data in this section was presented as tissue distribution extrapancreatic expression of insulin were presented as transcript per million. Graphic presentation was made using CIRCOS Circular Genome Data Visualization ^[29]^.

## RESULTS

Insulin is an important regulator of life and not surprisingly, is conserved throughout all animalia kingdom including animals from subphylum tunicata like *Ciona intestinalis*. Our phylogenetic analyze (Figure 1) demonstrates that insulin paralogues INS-IGF2, ins1 and insb have different origins, INS-IGF2 is an exclusive gene of Mammals except for murinae (mouse and rat) that do not have INS-IGF2 gene,but instead have ins1 gene. On the other and chinese hamster, that belongs to the same order (rodentia) of mouse and rat possess INS-IGF2 gene.

**Figure 1.**
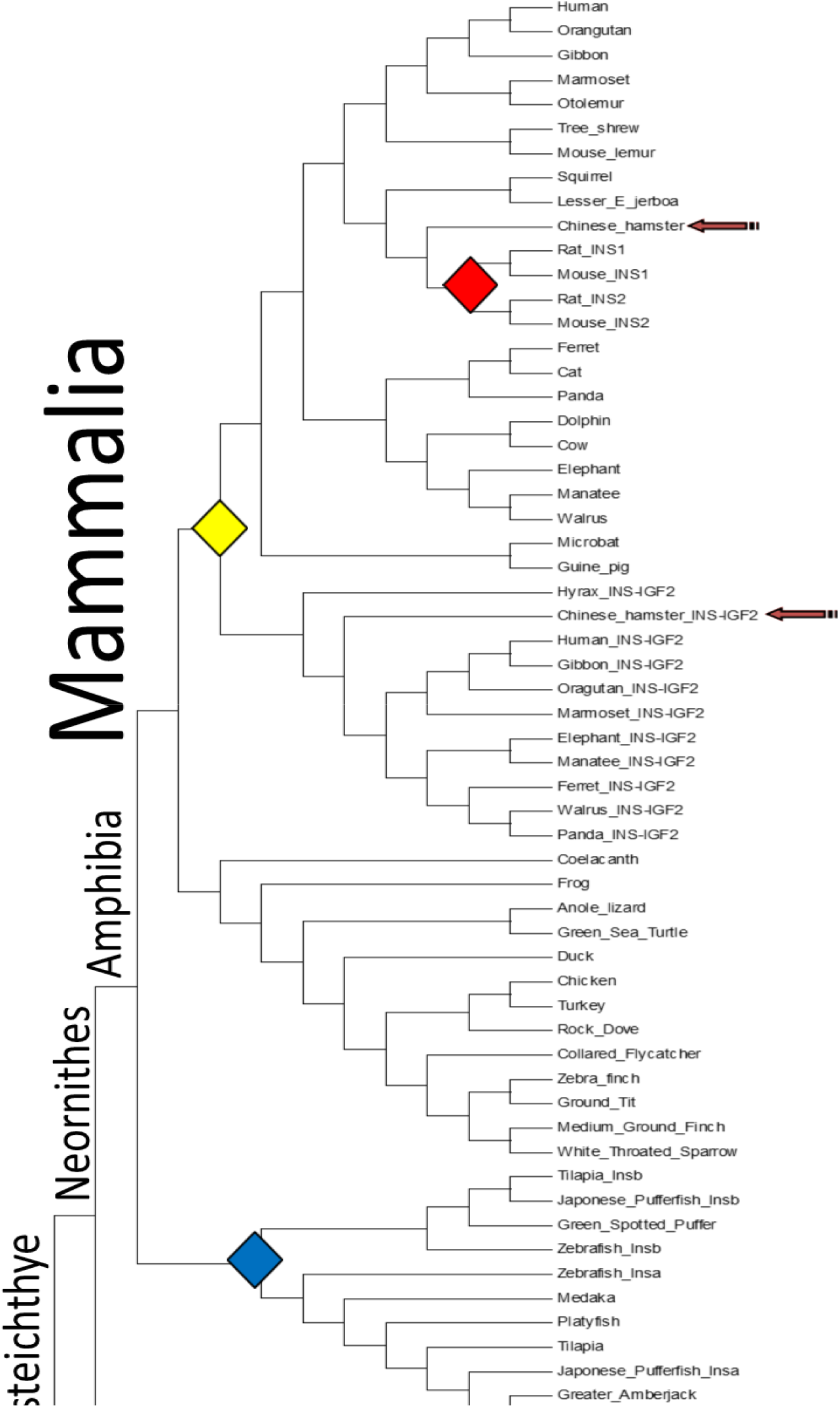
Phylogenetic tree of insulin, INS INS-IGF2, Ins2, Ins1, insa and insb proteins Sequences were obtained in Ensembl Genome Browser ^[20]^, aligned and phylogenetic and molecular evolutionary analyses were conducted using. For outgroup it was used insulin like 1 precursor protein sequence from *Ciona intestinalis*. Green circle INS, INS-IGF2 duplication, Blue circle Ins1, Ins2 duplication, red circle insa, insb duplication. Numbers at the branches represented the bootstrap support values.

The Insb paralogue seems to be exclusive of osteichthyes, and probably all species from this superclass have a second insulin gene, as we have identified possible regions for insb in the genome of stickleback and platyfish (data not shown). Transcript analyses (Figure 2) demonstrate that the regulatory elements of INS in human and ins2 in mouse are well conserved, since for each human transcript there is a similar mouse transcript (NP_000198.1 like NM_001185083.1, NP_001172026.1 like NM_008387.4, NP_001172027.1 similar to NM_001185084.1) except for ins2-006. Other transcripts are annotated in EGB ^[20]^, for ins2 (ins2-003, ins2-004 and ins2- 008), that predict protein structures smaller than the canonical sequence and this may need of further investigation to confirm it’s evidence. In zebrafish insa, presents only one known transcript, constituted of 3 exons like its human and mouse orthologues. Insb presents 3 different transcripts one with also 3 exons (NP_001034153.1) and two transcripts with 5 exons and a different coding DNA sequence, with the predicted protein being of more 49 aminoacids.

**Figure 2.**
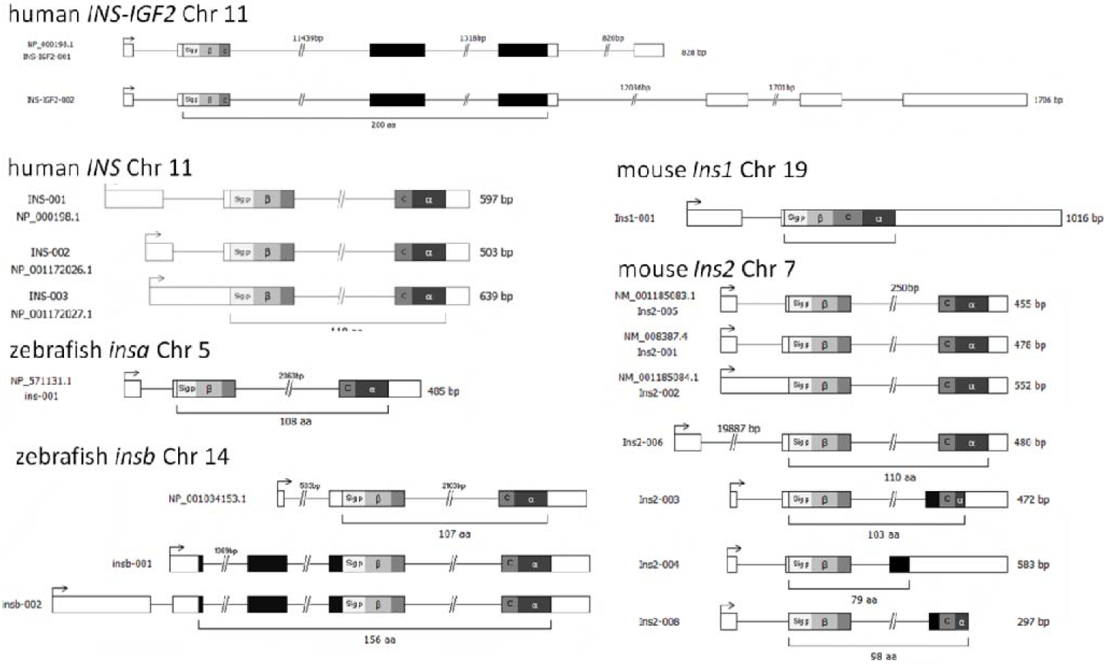
Transcripts for INS and INS-IGF2 in Human, ins2 and ins1 in mouse, insa and insb in zebrafish.

Two transcripts have been identified for INS-IGF2 were the two first exons share the same open reading frame (ORF) and coding DNA sequence (CDS) of INS transcripts, two other exons with CDS that encode for some unknown protein domain (confirmed by InterProScan ^[24]^, data not shown). Only one copy of ins1 is identified and this transcript presents a large size and structure similarities with of ins2, but without the second intron. Moreover, insa gene has only one known transcript and its structure is highly conserved compared with homologs INS and ins2, being identicaly constituted by 3 exons and with a similar size and structure. Insulin paralog insb of zebrafish presents 3 different transcripts, although the one annotated in NCBI (NP_001034153.1) seems to be a shorter version of the insb-001 and insb-002 present in EGB [20] and recently in our lab we have confirmed the existence of this two predicted transcripts by PCR amplification and sequencing. After, we aligned protein sequences (Figure 3) of human, mouse and zebrafish paralogs to understand the degree of conservation of the new insulin genes with ancestral genes. In human motif 1 and 2 of INS and INS-IGF2 are 100% identical as both proteins are translated from the same CDS, but INS-IGF2 lacks motif 3 signature from INS confirmed by InterProScan motif prediction [24]. On the other hand, mice paralogs have all three motif regions very well conserved, with an homology of motif 1 and 2 of 94,44% and motif 3 100%. Although not as conserved as mouse paralogs, in zebrafish, insb, seems to be reasonably conserved, as InterProScan [24] predicts the existence of the 3 insulin motifs, being the identity of insa and insb of 78,95% in motif 1 and 2 and of 40% in motif 3. In the past few years there has been a growing body of evidences showing that insulin can be expressed in several tissues beside pancreas. Here we show (Figure 4) data from transcriptome and microarray obtained in the repository data bases UniGene [25] and Expression atlas [28] demonstrating that in human, mouse and zebrafish there are indications of insulin expression in different tissues.

**Figure 3.**
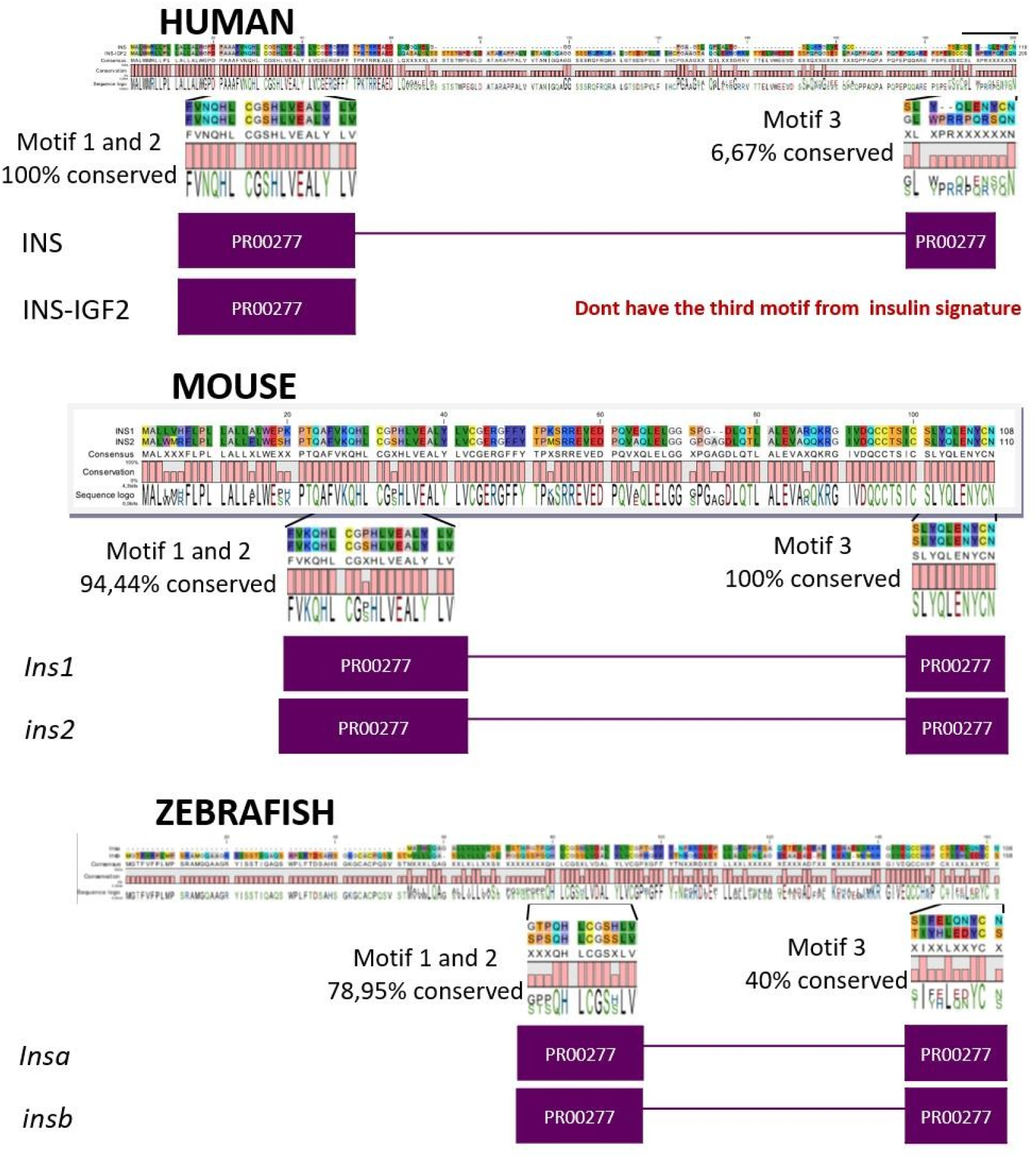
Mouse and zebrafish insulin paralogs are highly similar, but Human paralog INS- IGF2 don’t have the third motif of insulin protein.

**Figure 4.**
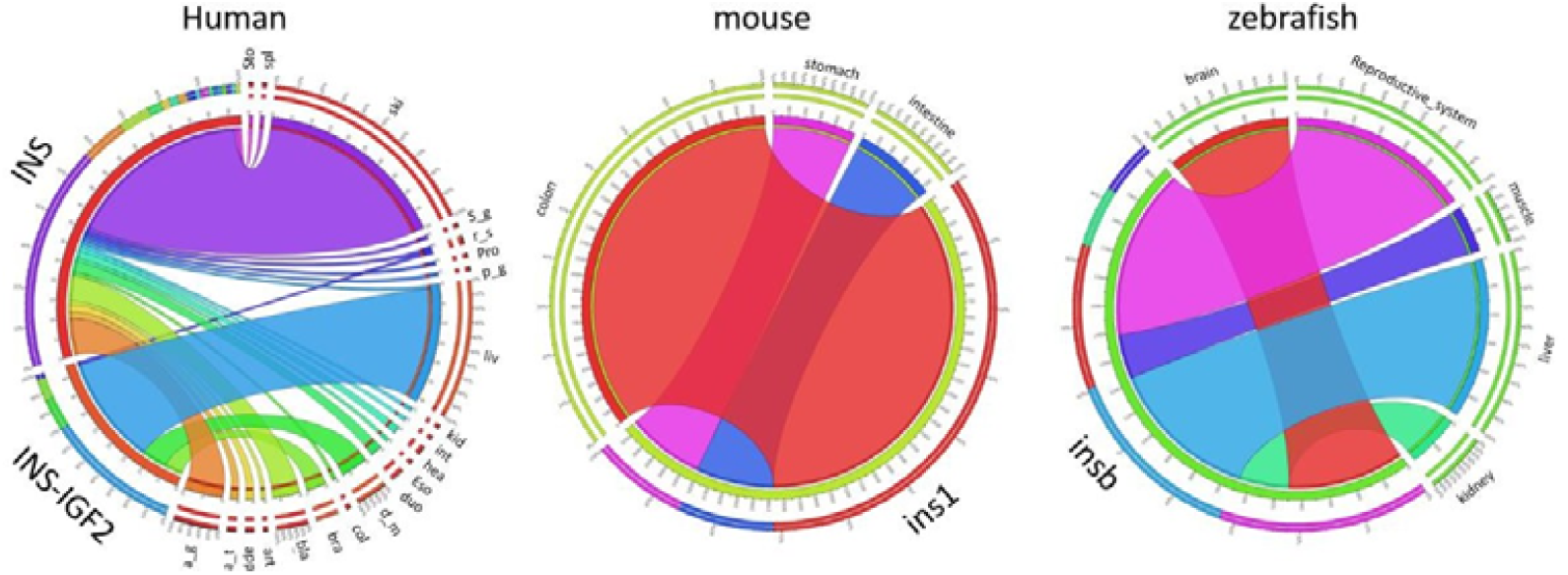
Gene expression data in different tissues, available in data banks, suggests that in all 3 species insulin is expressed outside the pancreas. In Human INS is expressed artery (art), kidney (kid), esophagus (eso), adipose tissue (a_t), adrenal gland (a_g), appendix (app), bladder (bla), muscle (mus), colon (col), duodenum (duo), heart (hea), liver (liv), prostate (pro), salivary gland (s_g), skin (ski), intestine (int), spleen (spl), stomach (sto), pituitary gland (p_g), reproductive system (r_s). INS-IGF2 was found in liver (liv), brain (bra), dura mater (d_m) and reproductive system (r_s). In mouse ins1 expression was found in colon (col), stomach (sto) and intestine (int), but no expression of ins2 was detected [27]. In zebrafish obtained data suggests that insa expression is restricted to pancreas while insb is expressed in brain, reproductive system, muscle, liver and kidney.

Human sequence analysis [26] showed that INS can be expressed in artery, kidney, esophagus adipose tissue, adrenal gland, appendix, bladder, muscle, colon, duodenum, heart, liver, prostate, salivary gland, skin, intestine, spleen, stomach, pituitary gland, reproductive system. Also, extrapancreatic expression of INS-IGF2 could be detected in liver, brain, dura mater and reproductive system [26]. In our review of mouse sequences, we could found evidence of expression in colon, stomach and intestine for the ins1 gene but no extrapancreatic expression of ins2 [27]. Similar results could be found in zebrafish as insa expression seems to be restricted to pancreas while insb was found in brain, reproductive system, muscle, liver and kidney.

## DISCUSSION

Strategies to overcome T1DM have been diverse, from vaccines ^[30]^, stem cells and pancreas transplantation ^[31]^, drug ^[32]^ and antibody therapy ^[33]^, with promising results in finding a possible cure. Other possible solutions, like extrapancreatic expression of insulin, have taken a marginal place although it could have some therapeutic potential in specific cases. The first question that we tried to answer was if INS-IGF2 could mimic the role of its ancestral. INS- IGF2 does not have all the conserved motifs of INS and although this may suggest other functions for this gene, data bases like STRING ^[34]^ recognize a protein interaction with INSR. INS, INS-IGF2 duplication event in mammals preceded the ins1 and ins2 duplication. It is possible that the conservation of ins1 led to the elimination of INS- IGF2 in Murinae, probably because it presents a protein structure extremely like ins2 and can, with accuracy, mimic ancestral gene function better than INS-IGF2. The cofunctionalization of ins1 and ins2 could be demonstrated in KO mice lacking ins2. These mutants did not present signs of diabetes and could stay normoglycemic by overexpressing ins1and increasing β-cell mass ^[9]^. Finally, ins1 replaced INS-IGF2 in Murinae genomes, because it was more efficient in performing the role ins2. Nevertheless, part of the INS-IGF2 protein sequence that is different from the INS, lacks any known domain that could also suggest neofunctiolization for this gene, although this has not yet been described. Additionally, a recent published work ^[35]^ observed that after trying to detect INS- IGF2 by both Western blotting and immunohistochemistry in a human beta cell line they only detected cross reaction to native proinsulin, concluding that there is no evidence of a mature protein from INS-IGF2. Further investigations must be performed to understand if INS-IGF2 can have some biological function or if it just a pseudogene. Evidence of extrapancreatic expression of insulin are becoming the rule rather than the exception. Large scale genome-wide association studies like The Genotype-Tissue Expression (GTEx) project ^[26]^, are enabling a global view of the gene expression in Human tissues. In this work the researchers could detect INS expression in twenty different tissues and INS-IGF2 in four. Transcriptome analyses in mouse and zebrafish suggest that extrapancreatic expression of insulin is achieved by ins1 and insb^[25,27]^. Although we look at this data with enthusiasm, we have the perfect conscience that the levels of expression, in some tissues, are very low and that proofs of protein secretion are needed. Evidence of extrapancreatic and extrathymic translation of insulin have been demonstrated in mice ^[16]^. Although in this case these insulin producing cells had origin in bone marrow and migrated to different parts of the body and were associated with the onset of diabetic neuropathy. Other studies could see advantages in the secretion of insulin by tear film cells, under diabetic conditions ^[18]^. Recently in our lab, we could detect both insa and insb expression in kidney, muscle and bone in WT zebrafish under nondiabetic conditions. Also, in a diabetic zebrafish model of diabetes, we could detect increased expression of both genes in bone after treating them with a vitamin D analog and a calcimimetic ^[36]^, demonstrating that not only pancreatic INS expression is sensitive to some types of treatments ^[37,38]^, but also that this drugs can have the same effect in other cell types including bone. In this work we have gathered evidence that in mice, and probably in zebrafish, insulin paralogues can regulate glucose homeostasis, but in Human the answer is not clear. Extrapancreatic expression of insulin occurs in the three studied species and, at least in zebrafish, it is sensitive to drug therapy. If confirmed, the therapeutic potential of this event would be of considerable relevance for the treatment of diabetes, as this could represent a simple, reliable and safe way for self-production of insulin.

## Acknowledgements

This study received Portuguese national funds from FCT - Foundation for Science and Technology through projects UIDB/04326/2020 (DOI:10.54499/UIDB/04326/2020), UIDP/04326/2020 (DOI:10.54499/UIDP/04326/2020) and LA/P/0101/2020 (DOI:10.54499/LA/P/0101/2020).

